# High-human acuity of speed asymmetry during walking

**DOI:** 10.1101/2020.10.28.359281

**Authors:** Pablo A. Iturralde, Marcela Gonzalez-Rubio, Gelsy Torres-Oviedo

## Abstract

Despite its central role in the proper functioning of the motor system, sensation has been less studied than motor output in sensorimotor adaptation paradigms. This deficit is probably due to the difficulty of measuring sensation: while motor output has easily observable consequences, sensation is by definition an internal variable of the motor system. In this study we asked how well can subjects estimate relevant environmental changes inducing motor adaptation. We addressed this question in the context of walking on a split-belt treadmill, which allows subjects to experience distinct belt speeds for each leg. We used a two-alternative forced-choice perceptual task (2AFC) in which subjects report which belt they thought to be moving slower. We characterized baseline accuracy in this task for healthy human subjects, and found 75% accuracy for 75 mm/s speed differences. Additionally, we used a drift-diffusion model of the task that could account for both accuracy and reaction times. We conclude that 2AFC tasks can be used to probe subjects’ estimates of the environment and that this approach opens an avenue for investigating perceptual deficits and its relation to motor impairments in clinical populations.

## 2 Introduction

Despite its central role in the proper functioning of the motor system, sensation has been less studied than motor output in sensorimotor adaptation paradigms. This deficit is probably due to the difficulty of measuring sensation: while motor output has easily observable consequences, sensation is by definition an internal variable of the motor system. For example, there exists abundant literature characterizing the adaptation of motor behavior evoked by split-belt walking (i.e., legs moving at different speeds) under a variety of conditions (e.g. Dietz et al., 1994; Torres-Oviedo and Bastian, 2010; Ogawa et al., 2014; Mawase et al., 2014), and for different populations (e.g. Reisman et al., 2007; Finley et al., 2015; Sombric et al., 2017), but the contribution of sensory information to this task is less understood. Psychophysical studies, in which subjects report what they perceive in response to a given physical stimuli, have been used to investigate the sensed speed differences that drive motor adaptation during split-belt treadmill walking (e.g. Jensen et al., 1998; Lauzière et al., 2014; Hoogkamer et al., 2015; Wutzke et al., 2015; Vazquez et al., 2015; Statton et al., 2018; Leech et al., 2018). However, the methodological approaches followed by these studies might be limited for the purpose of quantifying sensation.

Current literature on perception of speed differences fails to consider factors influencing people’s responses such. For example, people can report difference perception for the exact same sensory stimulus when probed multiple times (i.e., the probabilistic nature of perceptual responses). Similarly, the time it takes for people to generate a response varies depending on how confident the subject is (i.e. subject-specific confidence in their responses). These factors are are important for inferring human sensation from perceptual tests (Ehrenstein and Ehrenstein, 1999). During split-belt walking the external stimulus which needs to be sensed, which induces adaptation by disrupting the gait pattern, is the speed difference between the legs.

Previous reports have assessed people’s perception of belt speed differences through variants of yes/no tasks, in which individuals report when they perceive the belts to move at different speeds (i.e. yes, there is a speed difference) or the same speed (i.e. no, there is not a speed difference). Regardless of the sensory stimuli presentation modality, perceptual responses to yes/no tasks depend not only on subjects’ sensation, but also on subject-specific decision criteria to convert sensory information into a response (Ehrenstein and Ehrenstein, 1999). For example, some subjects are more likely to favor a response (e.g. “yes”) when in doubt. Resulting in different responses to identical sensory information for different subjects. Thus, the impact of subject-specific decision criteria in these yes/no assessments suggests that other perceptual methods, such as the two-alternative forced-choice task, might be preferable to study sensation.

The two-alternative-choice task (2AFC) is preferable to yes/no tasks because it allows us to access what people feel despite of their confidence on the stimuli that they are detecting. Notably, on a 2AFC task, subjects are asked to judge which of two stimuli contains a certain signal or satisfies a certain property. Importantly, subjects are forced to respond between the two alternatives, regardless of their confidence on the decision. One might consider that it is easier to indicate that something is happening (e.g., yes, there is a difference), rather than what is happening (e.g., left side moving slower than right one). However, the sensation thresholds determined through 2AFC tasks tend to be smaller in magnitude than those determined through yes/no tasks (Green and Swets, 1966). Consistently, studies using the two-alternative forced choice (2AFC) task that we propose have shown that individuals can detect external stimuli without explicit awareness of said stimuli (Goldstein, 2009; Ehrenstein and Ehrenstein, 1999). Thus, while the declarative aspect of our perceptual task is undeniable, the 2AFC task will give us a metric of at least partial sensory information available for sensori-motor recalibration.

Forcing a choice is a method to infer subjects’ sensations that would not get reported with other perceptual tests.Critically, 2AFC eliminates subject-specific differences in value between the alternative responses that exist for the yes/no task, making it simpler to obtain a quantification of sensor acuity. In other words, the alternatives in a yes/no task (i.e., “yes” or “no”) are not necessarily valued in the same way by every person. Thus, individuals may set personal decision thresholds adjusting for the value given to each of them. In contrast, in the 2AFC method the two alternatives are presented in a symmetric way, such that there is no additional value in one choice over the other. This makes 2AFC the preferred methodological approach to obtain a measure of sensory acuity (Green and Swets, 1966).

While detecting accuracy is considered to be the main outcome measure of perceptual tasks, reaction times are key in the process of accumulation of sensory evidence before making a decision in a discrimination task (Pardo-Vazquez et al., 2019; Henmon, 1911). Moreover, by using a 2AFC task we have access to information on subjects’ reaction times, which indicates the available sensory information when analyzed through an appropriate model of how choices are made. One such mechanistic model that can systematically explore both accuracy and reaction times in a discrimination task is the drift-diffusion model (e.g. Ratcliff, 1978; Bogacz et al., 2006; Gold and Shadlen, 2007). In the drift-diffusion model (DDM) for 2AFC an evidence variable is used to represent the accumulation of information, and a choice is made when one of two alternative choice barriers, corresponding to the two possible responses, is reached. Thus, the DDM offers a principled way to link accuracy and reaction times in the 2AFC as two expressions of the same mechanism for gathering sensory evidence to make a choice.

Here we rigorously characterized the human ability to detect differences in belt speeds on a split-belt treadmill. The focus of our study was to evaluate sensory information available to subjects, rather than their confidence levels or other decision criteria involved in eliciting responses. Consequently, we used a 2AFC task for our perceptual assessment. We present quantifications of accuracy, reaction times, and estimates of perceptual thresholds across subjects and stimuli magnitude. Further, we used a drift-diffusion model to gain insight into the processes underlying subjects choices and reaction times. Our results may be used as normative data to compare to other populations whose sensory acuity may differ or to assess changes in perception, for example, as a consequence of sensorimotor recalibration or lesions to the nervous system.

## 3 Methods

### 3.1 Data Collection

#### Participants

*N* = 9 healthy subjects (24.6 ± 3.7 y.o., 6 female) completed the protocol. All of the subjects were right-footed (self-reported leg used to kick a ball) and two of the subjects were left handed (self-reported). The protocol was approved by the University of Pittsburgh’s Internal Review Board (IRB) in accordance to the declaration of Helsinki.

#### Testing protocol

The overall protocol subjects experienced is depicted in Figure 1A. Throughout the whole protocol, participants walked on a split-belt treadmill with a mean speed between their legs of 1.05*m/s*. The subjects were initially familiarized with the perceptual task they were going to be performing throughout the protocol by performing 6 repetitions of the task while being provided with both visual (a live graphic of both belts’ speeds) and verbal feedback from the experimenters. This ensured subjects understood the task and mapping between their actions (key-presses) and the changes in the speed of the belts. Following familiarization, subjects performed 2 to 4 blocks of data collection (as time permitted). Each block consisted of interleaved tied-belt walking (i.e., both belts move at the same speed for 25 strides) and the perceptual tasks at regular intervals. The blocks had a single presentation of the non-zero stimulus sizes, defined as an imposed belt-speed difference (Δ*v* = *v*_*R*_ − *v*_*L*_) at the beginning of each perceptual trial, and two presentations of the null trials in pseudo-random order (the same order for all subjects). There was a total of 24 trials per block (see figure 1B), where the stimulus sizes consisted on any of the following values: 0 (null), ±10, ±25, ±50, ±75, ±100, ±125, ±150, ±200, ±250, ±300, and ±350 mm/s. The treadmill was only stopped between blocks, but not within them, and subjects were allowed to rest and move as desired during the breaks. Even blocks were mirror images of the odd blocks, so if the +250*mm/s* was presented first in the odd blocks, then the −250*mm/s* was presented first in the even blocks and so on. This ensured balancing of positive and negative perturbations to minimize experimentally-introduced biases in responses. We collected a total of *M* = 30 blocks from the *N* = 9 subjects, with two subjects completing just two blocks, two subjects completing 3, and the rest completing 4 blocks each.

**Figure 1:**
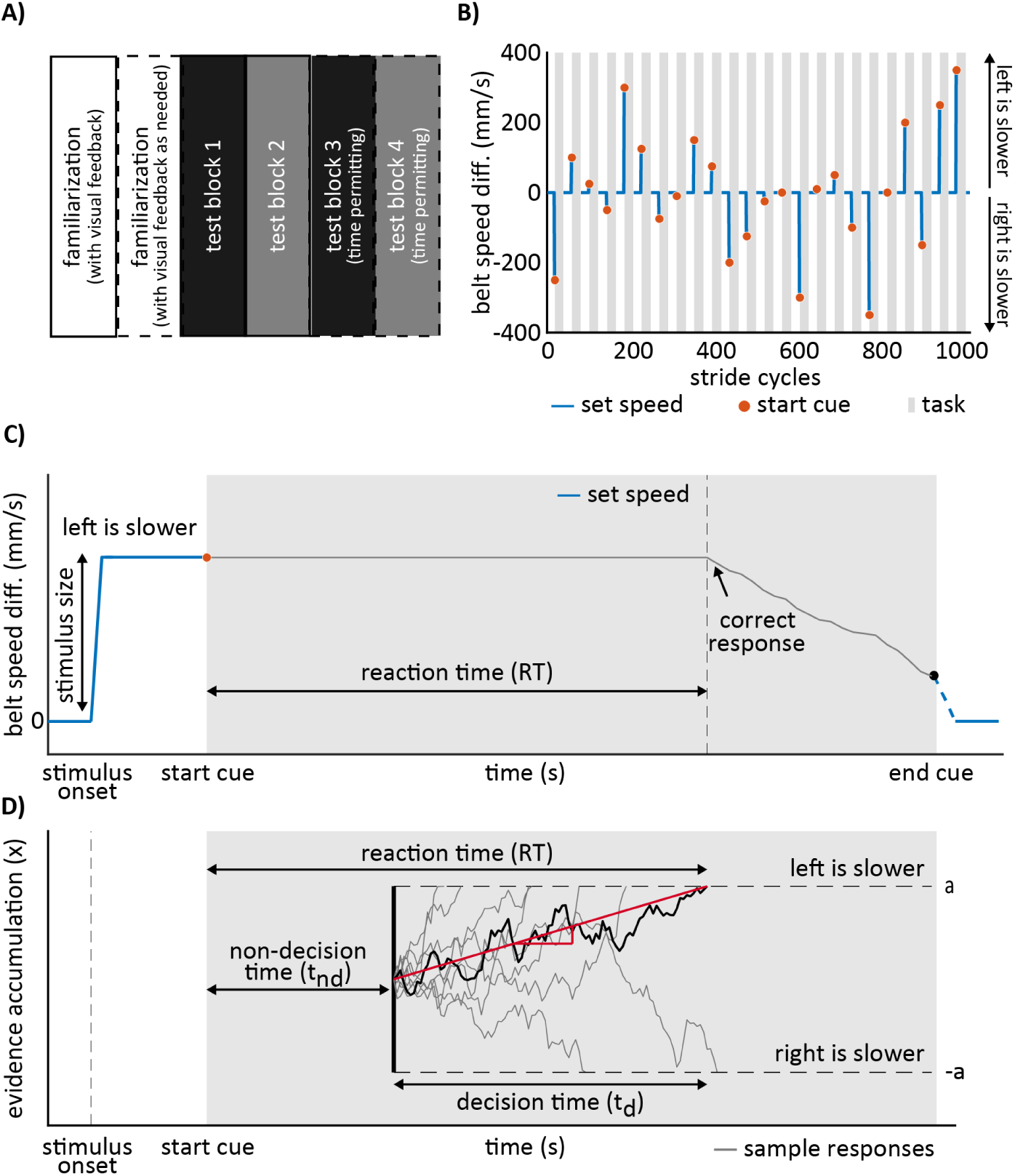
Protocol and methods for characterizing the perception of belt speed differences. **A)** Experimental protocol. Subjects completed one or two familiarization blocks followed by 2 to 4 testing blocks. **B)** speed difference profile for odd testing blocks. The profiles took the opposite values for the even blocks. Subjects performed a perceptual task in each of the shaded intervals. **C)** Perceptual task description. Tasks begun and ended with audio cues. Upon hearing the cue, subjects were instructed to identify the belt moving slower, and to make as many key-presses as necessary until the belts felt as moving at equal speeds. **D)** Drift-diffusion model schematic. The drift-diffusion model for 2AFC tasks represents the temporal evolution of a decision variable as a random-walk (black and gray jagged lines). Decisions are made when one of the two decision barriers is reached (dashed lines). Reaction time is composed of a non-decision and decision interval.

#### Description of the perceptual task

The perceptual task was designed to assess subjects’ perception of speed differences based on two methods: 2-alternative forced-choice followed by speed-matching (see Figure 1, panel C). The speed-matching component was a variant from previous perceptual tasks (Jensen et al., 1998; Vazquez et al., 2015) and is not analyzed in this study. Results from this portion of the task will be used in a future study comparing methods to track shifts in perception over motor adaptation. Every perceptual trial started with subjects walking with both belts moving at 1.05*m/s* followed by a sudden transition in belt-speeds to a speed difference whose value was unknown to the subjects (i.e., stimulus size). Subjects walked at this speed difference for a full stride cycle (i.e., time duration between two foot landings of the same leg) after which they heard an audio cue signaling the beginning of the response window (Figure 1C gray shaded area). Upon this audio cue, subjects had to press one of two keys (left or right) according to which belt they perceived to be moving slower. Response time (i.e., reaction time) and accuracy in the first key-press was used for the 2-alternative forced choice analysis. Subjects were also asked had to repeatedly press either key until the two belts felt like they were moving at the same speed (speed-matching component). At every key-press, the belt speed difference would change by either 6, 8 or 10 mm/s (equally probable, randomly chosen) such that the belt speed difference was either reduced (e.g., the subject pressed the key corresponding to the belt that was moving slower) or increased (e.g., the subject incorrectly pressed the key corresponding to the belt that was moving faster). Changes in belt speed were split symmetrically across the two belts such that the mean speed across the belts was constant throughout the experiment (1.05 m/s). The response window lasted for 24 strides, after which they would hear a second, different, audio cue indicating the end of the perceptual task. Subjects wore noise-cancelling headphones and a drape that blocked vision of their feet throughout the protocol to remove any additional auditory or visual information influencing the responses. The headphones were also used to provide the start/stop audio cues, as well as a clicking sound at each key-press to avoid any impression that the system may not be detecting their actions.

### 3.2 Perceptual Trial Exclusion

Trials where subjects did not respond and the first trial in each block were excluded from analysis. Non-response trials consisted of 1.1% of all recorded trials (8 out of 720), and occurred in small stimulus size trials only (i.e., 5/60 of ± 25 mm/s trials, 1/60 of ± 50 mm/s trials and 2/120 of ± 0 mm/s trials). We also eliminated the first perceptual trial for every block because we realized that other factors (e.g., distraction) beyond actual perception had an impact on the initial accuracy metric for every block. For example, subjects’ accuracy probed at 250 mm/s (which is a very salient speed difference) was much smaller when presented at the beginning than within the block (64% and 81% respectively).

### 3.3 Perceptual Task Outcome Variables

From each presentation of the perceptual task, we extracted two outcome variables that were subsequently used for analysis, both from the two-alternative forced-choice portion of the task: choice (first key pressed) and reaction time (time until first keypress). Results from the speed-matching portion of the task are not analyzed here. A ‘left’ choice was defined as a ‘left is slower than right’ response. A ‘right’ choice is defined analogously. Choices were converted to accuracy scores for some analyses. A response was considered accurate if the subjects’ choice indicated correctly the belt that moved slower. Accuracy was not defined for null trials (i.e. 0 mm/s stimulus size), as there is no correct choice. Reaction time was defined as the interval between the starting audio cue and the first keypress. Different from accuracy, reaction time is defined for all trials, including null ones.

### 3.4 Quantification of Perception Through the 2AFC Task

#### Perceptual thresholds (JNDs) and point of subjective equality (PSE)

We used the 2AFC task to estimate two important quantities in the characterization of perception: the just noticeable difference or differential threshold (JND), and the point of subjective equality (PSE). The JND or threshold is defined as the minimum magnitude difference between two stimuli that can be reliably perceived by the subject as distinct (e.g. Green and Swets, 1966). If we adopt a deterministic view of sensation, where a specific stimulus is either always detected or never detected, this definition matches the notion of a transition point between detection and non-detection. However, this all or nothing assumption rarely conforms to observations in perceptual studies, (Ehrenstein and Ehrenstein, 1999), requiring a probabilistic definition of thresholds. Thus we adopt the midpoint threshold definition (Goldstein, 2009), which is the stimulus value when probability of detection first exceeds the midpoint between its lowest and its largest possible values. In a 2AFC task such as the one used here, 75% accuracy is commonly used, given that subjects can be 50% accurate from merely guessing at the task. The point of subjective equality (PSE) is defined as the stimulus for which subjects choose equally between two options (50/50 choice ratio). This is a metric that quantifies potential biases in sensation (i.e., subjects are more likely to choose left than right or vice versa) and has been used in studies characterizing changes in perception following split-belt walking (Vazquez et al., 2015; Statton et al., 2018; Leech et al., 2018).

Both PSE and JND can be directly estimated by finding the stimuli values for which subjects select between the two alternatives with 50/50 and 75/25 ratios. While this is feasible in principle, accuracy estimates for each stimulus are very noisy, especially at the individual level were each stimulus was presented at most 4 times. Consequently, we decided against the direct approach and used responses for all stimuli presented to obtain a smooth estimate of the relationship between stimuli and responses in the task (i.e., psychometric function).

#### Psychometric curve fitting

As stated in the previous section, estimation of the PSE and JND was predicated on first characterizing the choice vs. stimulus size curve in this task. We did this through a maximum likelihood fit to the binary responses for both individual and group-averaged data. Specifically, we fit the binary left/right responses (not accuracy) data from each individual using a parametric logistic regression approach. The responses were modeled as coming from a binomial distribution with parameter *p*, where said parameter was taken to be a logistic function of a linear combination of the factors of interest (summarized in the function *µ*), as shown in Eq. 1.

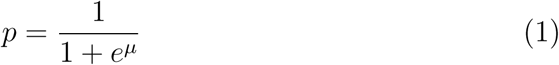

Initially, we considered two main factors that explain subject choices at each trial: a bias (intercept) term, and the stimulus size (Δ*V*_*k*_). We then considered the possibility that three exogenous factors may affect subject choice and consequently our JND estimate: task learning, habituation, and dominance. To control for these spurious effects we included three additional factors to our regression: previous stimulus size (Δ*V*_*k*−1_), a current stimulus by block number interaction (Δ*V*_*k*_ *× b*_*j*_, where *b*_*j*_ ∈ [0, 1, 2, 3] indicates the block number), and a term that depends solely on absolute stimulus size (|Δ*V*_*k*_|). This results in a five-factor model as shown in Eq. 2.

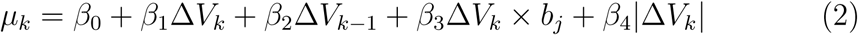

The intercept (*β*_0_) term quantifies a potential bias in responses. The second term (*β*_1_) quantifies sensitivity to stimulus size. The third term (*β*_2_) quantifies subjects’ habituation to previously presented belt speed differences. That is, because of prior experience subjects responses may be changing. In its simplest form, this can be taken as the responses to each stimulus being affected by the immediately preceding stimulus size. A negative value represents that having two stimuli of the same sign makes the second one more difficult to identify, while a positive value represents the opposite. The fourth term (*β*_3_) quantifies learning in the task, such that subjects would have a better performance in perceptual tasks presented later in the experiment. A significant value of *β*_3_ indicates a change in the slope of the psychometric functions for different blocks. A positive value would represent subjects getting increasingly better at the task (sharper transition between left and right choices) which is the expected effect, if any. Finally, the absolute stimulus size term (*β*_4_) would indicate a dominance effect. That is, it would indicate a higher sensitivity to one specific belt moving faster.

The model was first fit to all responses (pooled across subjects), to select the relevant factors among those considered. The model fitting was done using Matlab’s (The Mathworks, Inc., Natick, Massachussets, United States) *fitglm* function. A stepwise procedure was used to drop non-significant terms from the model one at a time. The criterion for dropping terms was a p-value larger than 0.05 for the likelihood ratio test of the model with and without the corresponding term, under a *χ*^2^ distribution with one degree of freedom. This procedure is implemented by Matlab’s *stepwiseglm* using the deviance criterion. Group regression results showed that only the bias and stimulus size factors (*β*_0_ and *β*_1_) were significant. Consequently, we fitted individual choice models considering those two factors only, as shown in Eq. 3.

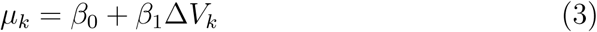

#### Estimation of the JND

Given the psychometric fits described above, it can be shown that for an unbiased subject 75% accuracy happens at stimuli values of 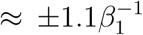. Consequently, we use the estimate 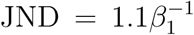. More generally, this JND quantification can be interpreted as the increase in stimulus needed to go from 50/50 response proportion to a 75/25 proportion (in favor of either response) if all other parameters and factors are held constant.

#### Estimation of the PSE

For the same psychometric fits described above, the belt speed difference for which subjects show equal proportion of responses is PSE = −*β*_0_*/β*_1_. We note that for healthy subjects the PSE in this task is expected to be close to 0 mm/s, such that if both belts are moving at the same speed, half of the time subjects will choose left and the other half right.

### 3.5 2AFC Decision Making as a Drift-Diffusion Process

The drift-diffusion model (DDM) is a model of decision making with noisy evidence. In this model, subjects are assumed to accumulate evidence in time until sufficient evidence is gathered and a decision is made. Here we consider the simplest form of DDMs for a two-alternative choice task. The evidence gathered up until time *t* is represented as the continuous variable *x*(*t*). Whenever *x*(*t*) goes above the barrier *a*(*t*) we say that a left choice has been made, and whenever it goes below *b*(*t*) we say a right choice has been made (see Figure 1D). Whenever the first barrier crossing happens, the trial terminates. We assume starting point is unbiased (i.e. that the starting point is equidistant to both decision thresholds, and thus subjects have no preference for either response), and fixed decision barriers in time. Thus, without loss of generality we take the starting point of the decision variable to be *x* = 0 and the barriers taken to lie symmetrically so *a*(*t*) = −*b*(*t*) = *a*. The simplest DDM can then be characterized by three additional parameters: the noise level (*σ*), the drift-rate (*r*), and the non-decision time (*t*_*nd*_). The model separates the evolution of *x*(*t*) into two stages. First there is a non-decision stage, representing a delay (*t*_*nd*_) in the beginning of the evidence gathering (Eq. 4).

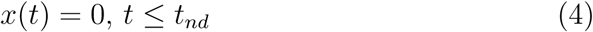

In the following stage the evidence gathering process is modeled as a continuous stochastic process with the following evolution (also known as a Wiener or Brownian motion process with drift):

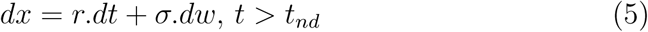

Where *r* and *σ* are constants (for a given stimulus), *dx* refers to the change in accumulated evidence for an infinitesimal time interval *dt*, and *dw* is a zero-mean normal process such that *dw* ∼ *N* (0, *dt*) for that same time interval. This equation can be interpreted as a process that accumulates evidence linearly (in time) through the given drift rate *r*, but is affected by additive noise also accumulated in time.

We note that despite there being four parameters for the model (*a, σ, r*, and *t*_*nd*_) the model is scale-invariant, such that proportionally scaling the values of *a, σ*, and *r* by the same amount results in the same predicted behavioral outcomes. Hence, without loss of generality we define “a” to be equal to 1 (Wagenmakers et al., 2007). Then the probability of the process hitting one particular decision barrier (e.g., the probability of a left choice being made, *P*_*L*_, which corresponds to the positive or upper decision barrier; Figure 1D), and the mean reaction time have closed-form expressions (Bogacz et al., 2006; Wagenmakers et al., 2007) given by:

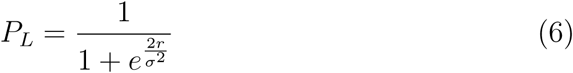

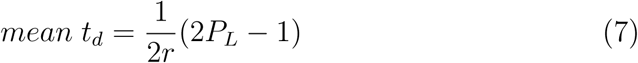

By extension, the probability of hitting the other barrier is *P*_*R*_ = 1 − *P*_*L*_. We note that the choice probability in the task if fully determined by *r* and *σ*, and satisfies 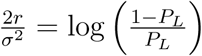.

Given the protocol design, where subjects experience the decision task for different stimulus sizes, it is necessary to establish the dependency of the model parameters (*r, σ, t*_*nd*_) to the different experimental conditions (i.e., different stimulus sizes). The experimental condition kept the mean speed as a fixed number 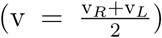. Therefore, it is possible to simplify the analysis on the dependency of the model parameters and the stimulus size such that the stimuli will be characterized jointly through their difference Δv = v_*R*_ − v_*L*_. Furthermore is is assumed that: 1) The non-decision time *t*_*nd*_ is independent of Δv. 2) Noise or diffusion rate is symmetric; that is, that a transposition of *v*_*R*_ and *v*_*L*_ leads to the same diffusion rate. The simplest such relation is to assume that the diffusion rate *σ* depends on the stimuli as *σ* = *σ*_0_ + *k*Δv^2^, where *σ*_0_ is a baseline noise and *k* scales the dependence on the stimuli. This relation can be interpreted as a second-order (Taylor) approximation of a more general relation that is symmetrical on the stimuli (i.e., that the diffusion rate is invariant to flipping the speeds of the two belts). 3) Finally, we assume that choice in the task must scale with stimulus size. Because choice is completely determined by 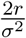, the simplest such relation is 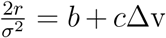, where *b* represents a bias term and *c* a scaling term. The drift rate *r* is then implicitly related to the stimuli. Similar to before, this can be interpreted as a first-order approximation of a more general dependency of 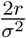 on Δv. We note that the model then has five scalar degrees of freedom (*b, c, t*_*nd*_, *σ*_0_, and *k*).

Using these definitions, the model was fit through a two-stage procedure: First, we find the maximum likelihood fit to choices (left/right), which determines the parameters *b* and *c* (Eq. 6). Second, we perform a least-squares fit to mean reaction times, which results in estimates of *t*_*nd*_, *σ*_0_ and *k* (Eq. 7). We will consider the special case with absence of signal-dependent noise (*k* = 0 and *σ* = *σ*_0_ is a constant), where the drift in the DDM is the only model parameter dependent on the stimulus size. Moreover, we analyzed the case in which *k* ≠ 0, where both the drift and the diffusion term in the DDM are dependent on the stimulus size (i.e., there is signal-dependent noise). Parameters were fit to each individual separately, but the resulting models are presented by averaging across all subjects for visualization purposes.

### 3.6 Data and Code Availability

Perceptual data and the code used for all analyses and creation of figures in this work are available at [THIS HAS BEEN ANONYMIZED, WILL REPLACE WITH PROPER URL AFTER MANUSCRIPT ACCEPTANCE]Kinematic and kinetic data from subjects while performing the task is available upon request.

## 4 Results

### 4.1 Characterization of Sensation Through the 2AFC Task

To answer the question of what factors could influence subjects’ perception of the speed at which their legs moved, subjects performed the 2 alternative forced choice task described in the methods. We fitted a logistic regression function to the averaged pooled responses (Figure 2A, black solid line), considering the effect of several factors beyond stimulus size. However, only stimulus size (*β*_1_ = 0.012 ± 0.001, mean ± standard error) significantly influenced subjects’ choices (*p* = 2.45×10^−34^). The intercept coefficient (*β*_0_ = 0.192 ± 0.098, mean ± standard error) almost reached significance (*p* = 0.051), indicating a potential group bias. However, this effect was mainly driven by a single subject with a large bias, and thus, should be interpreted carefully. This term is kept in the model for better fitting of the data on an individual level. Lastly, the other factors that were studied did not significantly affect subjects’ responses, such as leg dominance (*p* = 0.93), previous stimulus size (*p* = 0.25), or stimuli habituation (i.e., stimulus size by block number interaction, *p* = 0.17).

**Figure 2:**
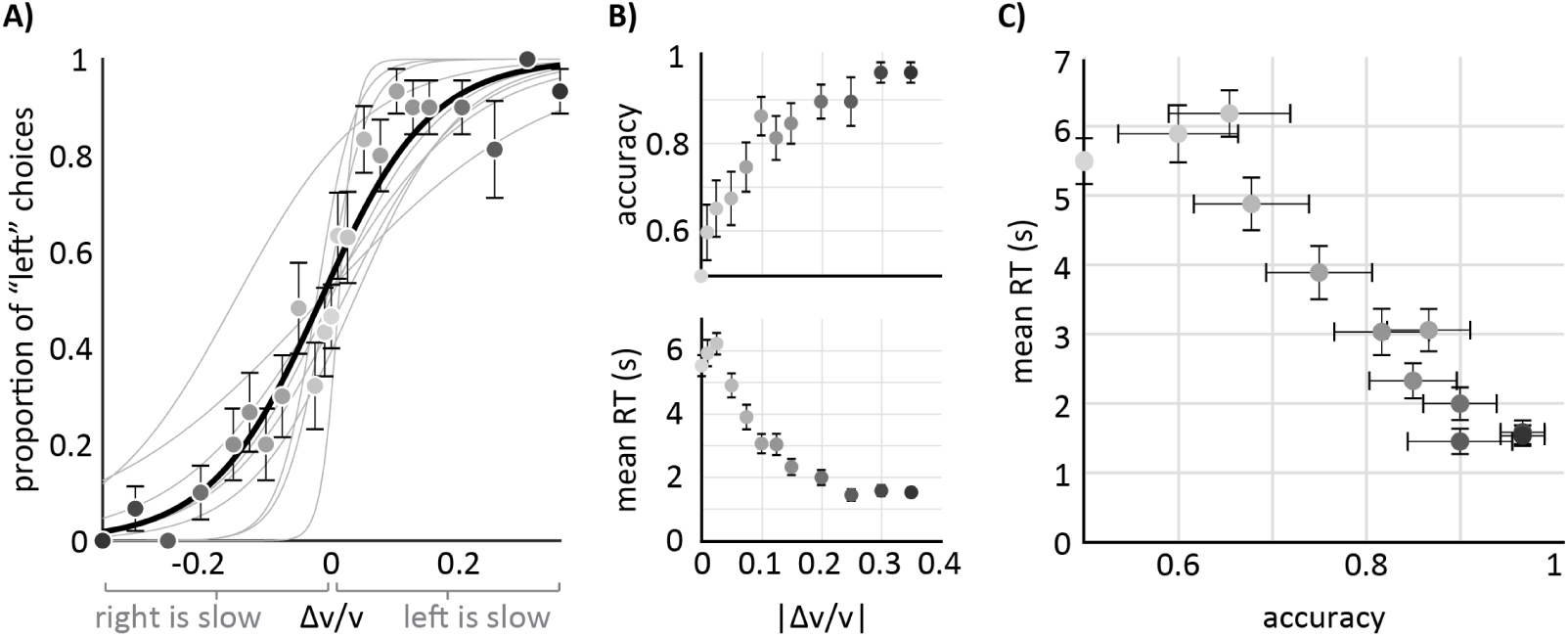
Choice, accuracy, and mean reaction times vs. stimulus size. **A)** choice as a function of stimulus size. Circles indicate group average responses across subjects (± standard error of the mean), thin gray lines represent logistic fits to individual data (see Methods), thick black line represents the average of the individual logistic fit curves (not the logistic fit to group averaged data). **B) Top:** accuracy as a function of absolute stimulus size. Both the data and model fits are the same as in the left panel, but averaged across positive and negative stimuli. Circles indicate experimental data (group average ± standard error). **Bottom:** mean reaction time as a function of absolute stimulus size. Circles indicate experimental data (group average ± standard error). **C)** mean reaction time vs. accuracy. Circles indicate experimental data (group average ± standard error). Note that the color gradient in the circles among B and C depend on the absolute stimulus size.

#### JND (group level)

The JND was estimated by smoothing available data and finding the point at which the logistic model, fitted to the subjects’ pooled data, crossed the 75% accuracy value. The group-level threshold (see methods) is 95 mm/s (equivalent to a 9.1% Weber fraction). The Weber fraction for this sensory modality is given by the threshold as a fraction of the mean belt-speed (i.e., 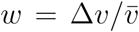). We note that the grouped data estimated over the 75 mm/s stimulus, has a group accuracy of exactly 75% (Figure 2B, top panel). We conclude that the perceptual threshold defined by 75% accuracy in the task is on average located approximately at 7% (Weber fraction) across the population tested.

#### Individual differences in accuracy, PSE, and JND

Individual subjects displayed a large range of behaviors in the 2AFC, with accuracy varying from 65% to almost 95% for individual subjects across all trials (see Figure 3A. Large reaction time differences were also observed, with mean values ranging from approximately 2.5 to 5.5 s. For PSE and the JND determination, choices were modeled for each individual considering the same two factors as in the group level (stimulus size and intercept, Figure 2A, gray lines). A representation of individual parameter estimates is shown in Figure 3 (Panel B: PSE; Panel C: JND). Only one of the subjects showed a significant bias (PSE, Subject 2), which we believe was driving the intercept term in the group level regression. A large range of values is observed for the JND estimate obtained from this model, implying that some subjects have much sharper discrimination curves than others. The observed threshold range was 18 to 191 mm/s, with an average of 96 ± 59 mm/s (mean ± standard deviation).

**Figure 3:**
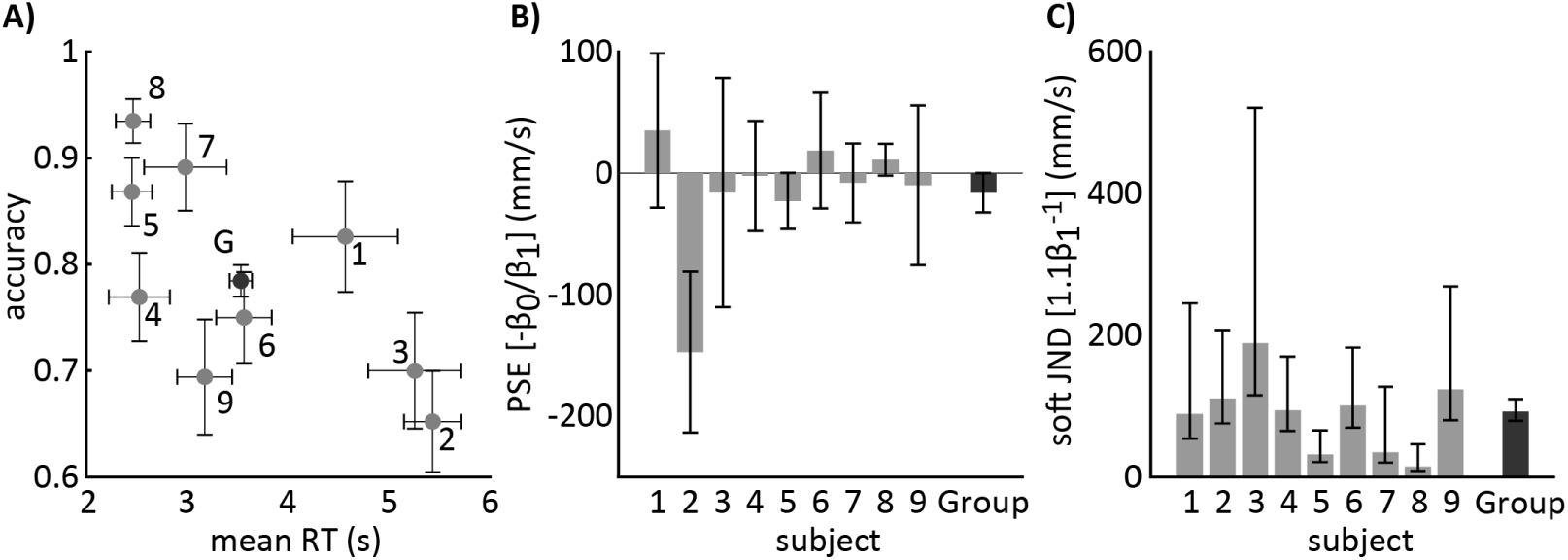
Individual variability in belt speed difference perception. **A)** mean reaction times (± standard error) vs. mean accuracy for each subject (numbered) and the whole Group across all stimulus sizes. **B)** estimates of point of subjective equality (PSE, thick bars) and 95% confidence interval (CI, errorbars) from logistic regression models. Best estimate and approximate confidence intervals are propagated from *β*_0_ and *β*_1_ estimates (PSE=−*β*_0_*/β*_1_) presuming fixed *β*_1_ to its maximum likelihood value. **C)** just noticeable difference (JND) estimate with 95% CI (errorbars). Values are computed as 1.1*/β*_1_. Confidence intervals are computed by applying the same transformation to the edges of the CI of *β*_1_. This results in skewed CIs.

#### The drift-diffusion model can explain accuracy and reaction times in this task

This model represents the decision-making process in the 2AFC task as the random walk of a variable quantifying the evidence accumulated over time. A choice is made once the evidence exceeds some pre-determined values (see Figure 4, top). The drift-diffusion model (DDM) could adequately fit experimental results. Figure 4 shows group-averaged data along with group-averaged model predictions. Model parameters were fit to each individual separately, assuming a linear relation between stimuli (belt speed difference) and drift rate.

**Figure 4:**
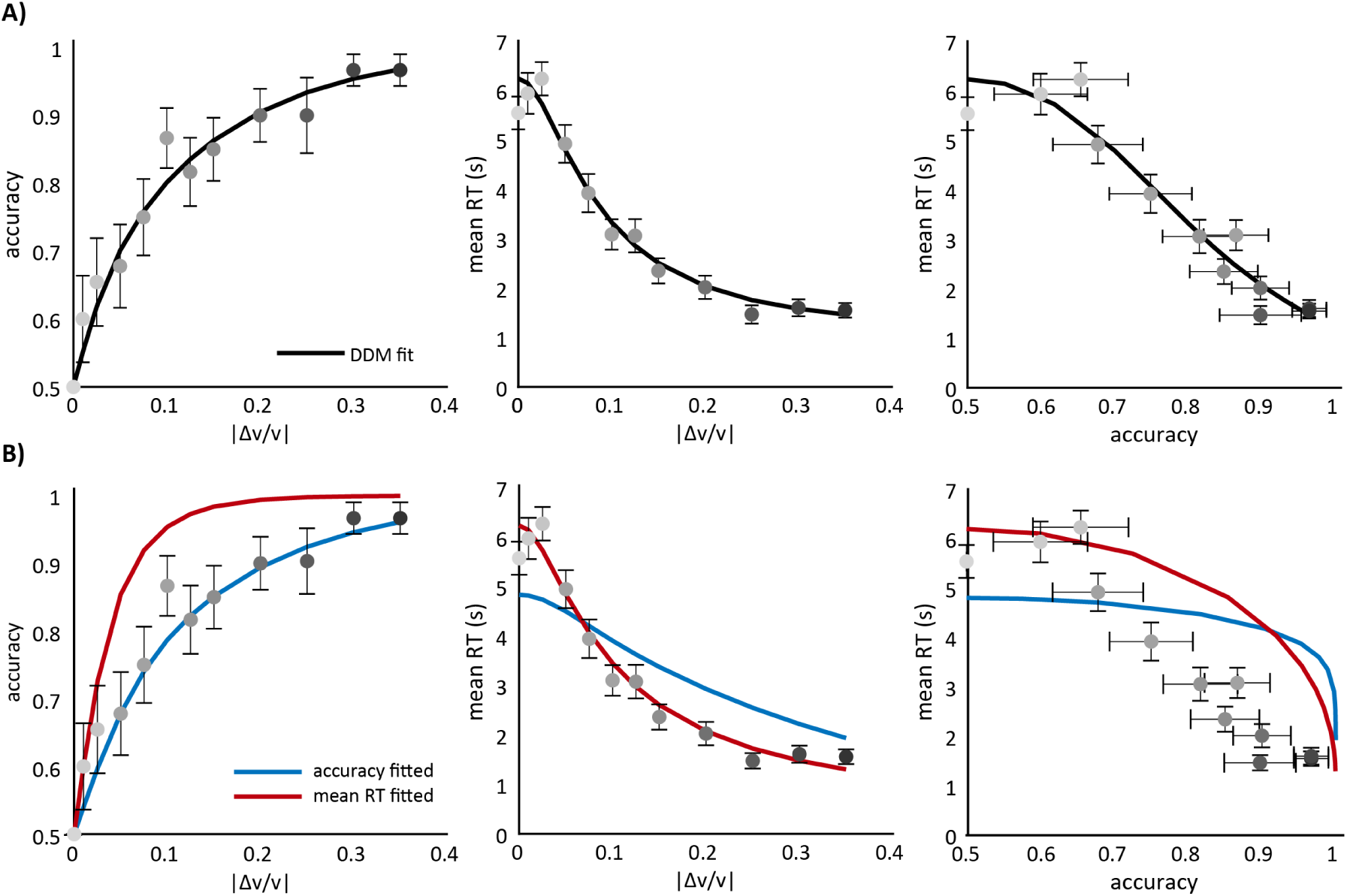
Drift diffusion models with fixed noise are insufficient to fit 2AFC task data. **A) Left panel:** accuracy vs. absolute stimulus size. Black line represents the model fit, circles indicate experimental data (group average ± standard error). **Middle Panel:** mean reaction time (RT) vs. absolute stimulus size. **Right Panel:** mean RT vs. accuracy. **B)** The model with fixed noise could be fitted to adequately explain accuracy (Blue line) or mean reaction time (Red line). **Left Panel:** accuracy vs. absolute stimulus size. **Middle Panel:** mean reaction time (RT) vs. absolute stimulus size. **Right Panel:** mean RT vs. accuracy.

We note that a DDM with fixed noise could be fitted to adequately explain accuracy or reaction times (RT), but it was not possible to describe both outcomes simultaneously (results not shown). For the parameters that best fit RT, the expected subject accuracy was higher than the empirical one. Conversely for the parameters that best fit accuracy the expected RT curve was flatter than the observations. We conclude that the DDM as presented is sufficient to characterize both subjects’ accuracy and reaction times if the model’s noise, or diffusion rate, is allowed to be stimulus-dependent.

## 5 Discussion

### 5.1 Quantification of Belt Speed Difference Sensation Through a 2AFC Task

This work presents a rigorous assessment of subjects’ sensation of differences in belt speeds on a split-belt treadmill. Specifically, we present results on the accuracy of subjects in identifying which belt was moving faster than the other as a function of the magnitude of belt speed differences. Different from prior studies on the topic that relied on variants of a yes/no task (Lauzière et al., 2014; Hoogkamer et al., 2015; Wutzke et al., 2015), we utilized a two-alternative forced-choice (2AFC). Yes/no task responses depend on both available sensory information and of subject’s valuation of the two alternative responses. For example, when asked if belts are moving at the same speed, subjects favor one response when in doubt, while others may prefer the alternative answer. However, none of the prior studies explicitly account for this decision criteria. The 2AFC presents alternative responses in a symmetric design, so differences in value between the alternatives can be assumed to be nonexistent. This allows for inference about the sensory information used to arrive at a response without needing to consider subjects’ decision criteria, making 2AFC the preferred task when the objective is to assess sensory information and processes (Green and Swets, 1966). Consequently, we believe the results from our work represent the most accurate report to date on healthy young persons’ sensation on this task.

### 5.2 Middle Point JNDs are a More Meaningful Metric of Sensitivity than Null Hypothesis JNDs

The just noticeable difference (JND) is a useful metric to summarize sensation of a particular physical stimulus. Three prior reports on sensation of belt speed differences in split-belt walking explicitly estimate JNDs in this context (Lauzière et al., 2014; Hoogkamer et al., 2015; Wutzke et al., 2015). However, these studies are not always explicit about what they define as the JND, which makes interpretation of experimental results harder. In this work we adopted and quantified our results through one commonly used definition: the middle point threshold. An alternative definition is the one given by the null hypothesis threshold (Goldstein, 2009). Both definitions (middle point threshold and null hypothesis) quantify different notions of what a JND is, and either or both may be adopted to characterize probabilistic relation between detection and stimulus size. However, we believe that the middle point JND quantification is more meaningful when characterizing sensitivity.

The null hypothesis JND can be qualitatively defined as the point below which stimuli are not reflected in sensory information that is available to subjects’ response mechanisms. Consequently, belt speed differences below this threshold should result in chance-level (50%) choices. Here we found that subjects, when taken as a group, were able to detect above chance levels, belt speed differences as small as 25 mm/s when the mean belt speed was 1.05 m/s. This corresponds to a 2.4% Weber fraction. Subjects’ accuracy in determining belt speed differences of 10 mm/s (the smallest tested here) was 60%, but this value was not significantly different from chance. These results suggest that the null hypothesis threshold in this context is certainly below 2.4%, and possibly lies between 1 and 2.4%.

While null hypothesis JND may make intuitive sense, it is impossible to positively prove its existence. Any negative findings (i.e., chance-level accuracy for some non-zero stimuli) can be explained away as a lack of statistical power to assess true accuracy. In our case, if the true underlying accuracy for 10 mm/s differences is 60% (as estimated here), then experimental determination of this value to be above chance levels with 80% power and a 5% type I error rate would require over 150 samples. Hence, one possible interpretation of our results is that the experiment was underpowered to detect a 60% accuracy rate for 10 mm/s differences in belt speeds. Of course, this problem can be controlled if a definition exists of what is the minimum accuracy level that we deem to be meaningfully above chance (e.g., defining that accuracy below 55 % is not meaningfully above chance even if it may be statistically significant for a large enough sample size). Thus, we believe estimation of accuracy rates for any particular stimulus size (e.g., defining estimates and confidence intervals of accuracy for any given belt speed difference) is a more meaningful way to describe sensation than ascertaining the existence of a precise cutoff point.

The middle point JND corresponds to the point at which subjects are more likely than not to correctly identify belt speed differences. In our task this point corresponds naturally to the 75% accuracy point in the accuracy vs. stimulus strength. This definition has the advantage of being a good descriptor of perceptual sensitivity, regardless of whether thresholds in the null hypothesis sense exist. Using this metric we found a threshold of 9.1% for group-averaged accuracy data. Further, we were able to quantify this for individual subjects, and we found a wide range of sensitivities at the individual level, with a population average also of 9.1% ± 5.6 % (mean ± standard deviation). Although the results from the psychometric fits suggest a threshold of 9.1%, the grouped data suggests a lower threshold (Figure 2B, top panel). Note that the grouped data estimated over the 75 mm/s stimulus has an accuracy of exactly 75%, suggesting that the group-level JND is at most 75 mm/s and the psychometric fit is overestimating the threshold. Notably, these estimates are well below previously reported perceptual belt speed thresholds.

Because of differences in threshold definitions, along with the previously mentioned methodological differences, comparisons between studies should be made cautiously. A summary of prior reports on split-belt treadmill JNDs is given in Table 1. Wutzke et al. (2015) explicitly use a middle point definition. The two other reports (Lauzière et al., 2014; Hoogkamer et al., 2015) implicitly adopt a non-probabilistic approach, quantifying thresholds from a single, or at most two, independent measurements. This approach appears more consistent with a null hypothesis framework, in which stimuli fall in either the undetectable (chance level accuracy) or the detectable (100% accuracy) categories. Whatever the definition, our results reflect higher sensory acuity from healthy subjects than had previously been implied. This is consistent with prior observations that subjects require less stimulus information to make a decision in forced-choice tasks (Ehrenstein and Ehrenstein, 1999).

Throughout this report we have presented metrics of sensitivity to belt speed differences as a Weber fraction, or % of mean belt speed. This normalization procedure is based on the finding, across several sensory modalities, that the ability to perceive a difference between two stimuli scales linearly with stimulus magnitude (Goldstein, 2009). This notion was not directly tested here, but prior reports on this context offer support for this normalization. Specifically, thresholds corresponding to equivalent Weber fractions have been found in studies using varied walking speeds (Lauzière et al., 2014). Verification of this relation is left for future studies.

**Table 1:** Summary of split-belt perceptual thresholds reported in the literature, presented as a fraction of mean belt speed as a normalization procedure (Weber fraction, 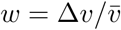).

| Authors | Population | Procedure | JND |
| --- | --- | --- | --- |
| Lauzière et al. (2014) | healthy elderly | y/n ascending limit | 12.8% |
| Lauzière et al. (2014) | healthy elderly | y/n descending limit | 16.2% |
| Wutzke et al. (2015) | chronic poststroke | y/n staircase | 26% |
| Hoogkamer et al. (2015) | healthy young | y/n ascending limit | 13% |
| Hoogkamer et al. (2015) | cerebellar | y/n ascending limit | 17.6% |
| this study | healthy young | 2AFC | < 9% |

### 5.3 The Drift-Diffusion Model is Sufficient to Describe Choices and Reaction Times in the 2AFC Task

Drift-diffusion models (DDMs) have been successfully used to describe a wide range of perceptual tasks (Ratcliff, 1978; Gold and Shadlen, 2007). One of the appeals of DDMs is that it offers a framework to relate perceptual outcomes (choice) with reaction times, which are not clearly related to the task objective, through a computational description that depends on only a few parameters. Further, the DDM can be interpreted as a formalization of a process of statistical inference based on sequentially acquired information Bogacz et al. (2006); Gold and Shadlen (2007). In our task DDMs with noise levels that are stimulus-dependent were able to describe both accuracy or reaction times as a function of stimulus strength. This validates the use of DDMs in this type of perceptual task, which differs from prior applications in that it is a movement task, and in which decisions are made over seconds to tens of seconds, rather than in hundreds of milliseconds.

Recent work has suggested (Pardo-Vazquez et al., 2019) that this type of decision-making models, when combined with experimental results made under different stimuli combinations, can be informative about the neural coding of said stimuli. Thus, it is of interest to understand if relations such as Weber’s law hold in this context too.

Relatedly, it would be interesting to study the problem of optimal decision barriers on this task as a function of a time vs. accuracy trade-off across different stimuli. The study of these potential modifications to the model is left for future studies.

